# Highly efficient SARS-CoV-2 infection of human cardiomyocytes: spike protein-mediated cell fusion and its inhibition

**DOI:** 10.1101/2021.07.30.454437

**Authors:** Chanakha K. Navaratnarajah, David R. Pease, Peter Halfmann, Biruhalem Taye, Alison Barkhymer, Kyle G. Howell, Jon E. Charlesworth, Trace A. Christensen, Yoshihiro Kawaoka, Roberto Cattaneo, Jay W. Schneider, Wanek Family Program for HLHS-Stem Cell Pipeline, Timothy J. Nelson (Director), Boyd Rasmussen and Frank J. Secreto

## Abstract

Severe cardiovascular complications can occur in coronavirus disease of 2019 (COVID-19) patients. Cardiac damage is attributed mostly to a bystander effect: the aberrant host response to acute respiratory infection. However, direct infection of cardiac tissue by severe acute respiratory syndrome coronavirus 2 (SARS-CoV-2) also occurs. We examined here the cardiac tropism of SARS-CoV-2 in human induced pluripotent stem cell-derived cardiomyocytes (hiPSC-CM) that beat spontaneously. These cardiomyocytes express the angiotensin I converting-enzyme 2 (ACE2) receptor and a subset of the proteases that mediate spike protein cleavage in the lungs, but not transmembrane protease serine 2 (TMPRSS2). Nevertheless, SARS-CoV-2 infection was productive: viral transcripts accounted for about 88% of total mRNA. In the cytoplasm of infected hiPSC-CM, smooth walled exocytic vesicles contained numerous 65-90 nm particles with typical ribonucleocapsid structures, and virus-like particles with knob-like spikes covered the cell surface. To better understand the mechanisms of SARS-CoV-2 spread in hiPSC-CM we engineered an expression vector coding for the spike protein with a monomeric emerald-green fluorescent protein fused to its cytoplasmic tail (S-mEm). Proteolytic processing of S-mEm and the parental spike were equivalent. Live cell imaging tracked spread of S-mEm signal from cell to cell and documented formation of syncytia. A cell-permeable, peptide-based molecule that blocks the catalytic site of furin abolished cell fusion. A spike mutant with the single amino acid change R682S that inactivates the furin cleavage site was fusion inactive. Thus, SARS-CoV-2 can replicate efficiently in hiPSC-CM and furin activation of its spike protein is required for fusion-based cytopathology. This hiPSC-CM platform provides an opportunity for target-based drug discovery in cardiac COVID-19.

**Author Summary:** It is unclear whether the cardiac complications frequently observed in COVID-19 patients are due exclusively to systemic inflammation and thrombosis. Viral replication has occasionally been confirmed in cardiac tissue, but rigorous analyses are restricted to rare autopsy materials. Moreover, there are few animal models to study cardiovascular complications of coronavirus infections. To overcome these limitations, we developed an *in vitro* model of SARS-CoV-2 spread in induced pluripotent stem cell-derived cardiomyocytes. In these cells, infection is highly productive: viral transcription levels exceed those documented in permissive transformed cell lines. To better understand the mechanisms of SARS-CoV-2 spread we expressed a fluorescent version of its spike protein that allowed to characterize a fusion-based cytopathic effect. A mutant of the spike protein with a single amino acid mutation in the furin cleavage site lost cytopathic function. The spike protein of the Middle East Respiratory Syndrome (MERS) coronavirus drove cardiomyocyte fusion with slow kinetics, whereas the spike proteins of SARS-CoV and the respiratory coronavirus 229E were inactive. These fusion activities correlated with the level of cardiovascular complications observed in infections with the respective viruses. These data indicate that SARS-CoV-2 has the potential to cause cardiac damage by fusing cardiomyocytes.

## Introduction

Severe acute respiratory syndrome coronavirus 2 (SARS-CoV-2), the coronavirus family member that most recently adapted to humans, is the etiologic agent of the coronavirus disease of 2019 (COVID-19). While the four coronaviruses endemic to humans (229E, NL63, OC43 and HKU1) impact mainly the respiratory tract and usually cause mild symptoms, SARS-CoV-2, as the other emerging coronaviruses SARS-CoV and Middle East Respiratory Syndrome (MERS), can cause lethal systemic symptoms [1].

Systemic symptoms caused by the three emerging coronaviruses include cardiovascular complications. In particular, SARS-CoV-2 infection causes myocardial disease in a significant fraction of COVID-19 patients [2]. Complications include worsening of pre-existing conditions and the onset of new disorders [3-5]. New disorders range from myocardial injury with or without classic coronary occlusion to arrhythmias and heart failure [3, 6].

Many cardiac symptoms have been tentatively attributed to aberrant host responses to acute respiratory infection [7, 8], but the complex mechanisms of cardiac disease are incompletely understood. As SARS-CoV-2 nucleic acids and proteins have been occasionally detected in cardiac tissue [9-17], productive SARS-CoV-2 infection of cardiomyocytes may directly cause disease. However, this hypothesis is difficult to verify experimentally. Rigorous analyses of cardiac tissue are restricted to rare autopsy materials, and there are few animal models to study cardiovascular complications of any coronavirus infection [1, 18].

To overcome these limitations, human induced pluripotent stem cell-derived cardiomyocytes (hiPSC-CM) have been used to model SARS-CoV-2 spread in cardiac tissue [19-23]. Focusing on hiPSC-CM from a developmental stage with peak expression of the SARS-CoV-2 receptor, angiotensin I converting-enzyme 2 (ACE2), we have established a new model of SARS-CoV-2 infection. In this model SARS-CoV-2 replicates more efficiently than in the hiPSC-CM models previously used. Electron microscopy analyses document large amounts of coronavirus particles both within exocytic vesicles and at the surface of infected cells, that form syncytia. By expressing the spike proteins of SARS-CoV-2 and other coronaviruses, we have gained insights into the mechanisms of their functional activation and which proteins have the potential to cause cardiac damage.

## Results

### Expression of virus entry factors in cardiomyocytes

We assessed whether ACE2, the SARS-CoV-2 receptor, and the spike-activating proteases TMPRSS2 and cathepsin B (CTSB) are expressed during the differentiation process of human embryonic stem cells into cardiomyocytes. The ACE2 transcription level peaked at day-20, those of cathepsin B remained stable, and TMPRSS2 transcripts were never detectable (**S1** Table). Thus, we characterized our hiPSC-CM at this differentiation stage. Super resolution immunofluorescence confocal microscopy documented cell surface expression of ACE2, and the striated F-actin organization typical of cardiomyocytes (Fig. **1A**). In particular, ACE2 receptors clustered in raft-like puncta diffusely distributed across the sarcolemma and extended into filopodia contacting adjacent cardiomyocytes (Fig. **1A**, arrow highlights filopodia).

**Fig. 1:**
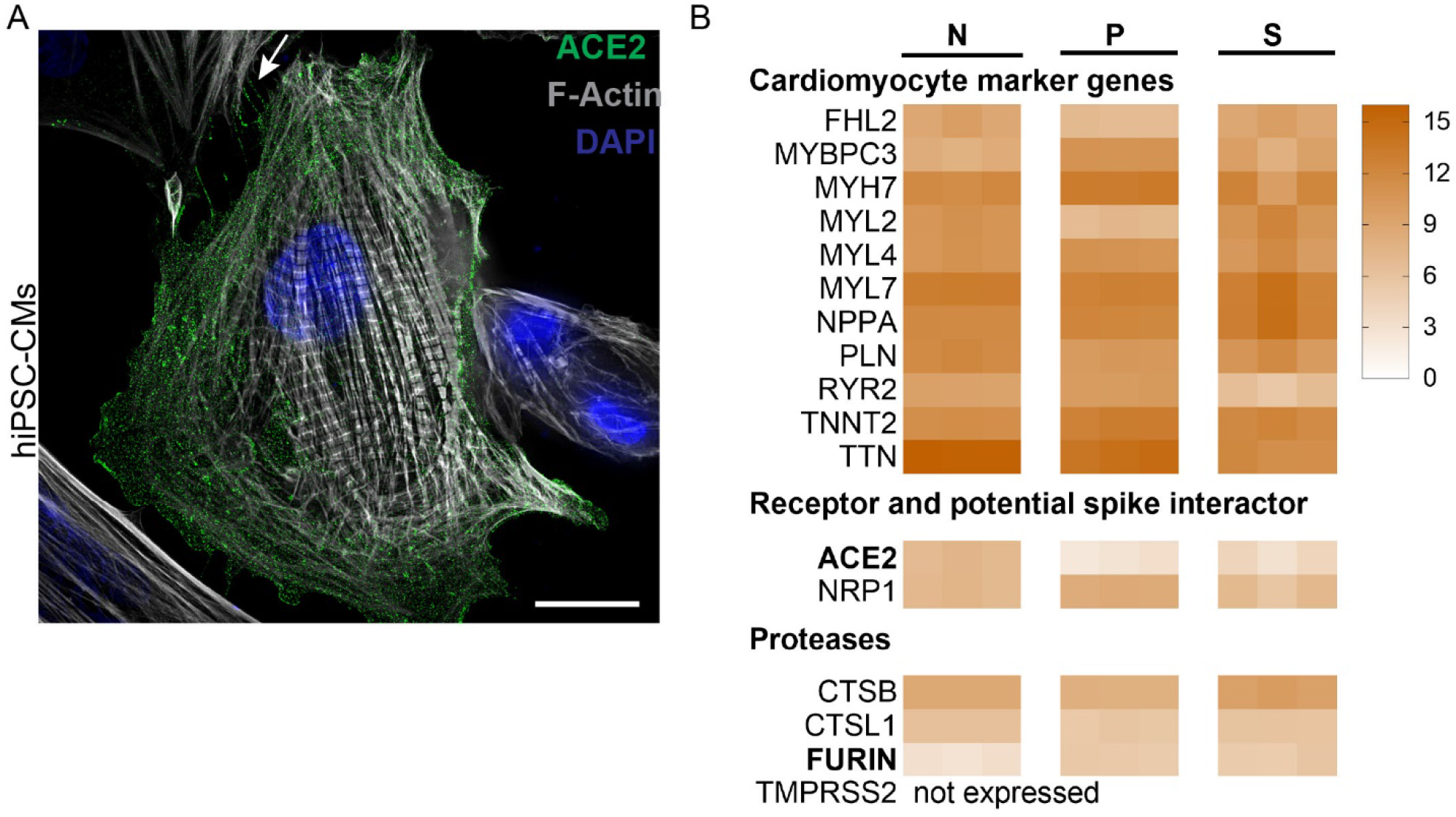
hiPSC-CMs express virus entry factors. (**A**) IF super resolution confocal microscopy analysis of ACE2 and F-actin expression in hiPSC-CMs. Scale bar, 10 µm. (**B**) Transcript levels of cardiomyocytes marker genes and of virus entry factors. Scale on the right: log2 trimmed mean of M values (TMM) normalized sequence read counts. N=this study; P=Perez-Bermejo et al., 2021 [21]; S=Sharma et al., 2020 [19]. Gene abbreviations: FHL2=four and a half LIM domains 2; MYBPC3=myosin binding protein C; MYH=myosin, heavy chain; MYL= myosin, light chain; NPPA=natriuretic peptide A; PLN=phospholamban; RYR2=ryanodine receptor 2; TNNT2=troponin T type 2; TTN=titin; ACE2=angiotensin I converting enzyme; NRP1=neuropilin 1, a potential spike interactor [53, 54]; CTSB=cathepsin B; CTSL1=cathepsin L1; TMPRSS2= transmembrane serine protease 2.

To further characterize day-20 differentiated cardiomyocytes, we analyzed their total cellular transcriptome by RNAseq. Cardiomyocyte differentiation markers were expressed in our hiPSC-CM at levels similar to those documented in two other hiPSC-CM lines used for SARS-CoV-2 infection studies (Fig. **1B**, compare N with P and S panels). The ACE2 receptor was expressed at higher levels in our cardiomyocytes than in those used in the other studies. In all three studies transcripts of the proteases cathepsins B, cathepsin L, and furin, were detected, but transcripts of the protease (TMPRSS2) that enables endosome independent viral entry in the lungs [24], were below detection levels (less than 0.5 counts/million in at least 2 samples).

### Highly productive cardiomyocyte infection

We inoculated two independent lines of spontaneously beating hiPSC-CMs with SARS-CoV-2 at 0.01 multiplicity of infection (MOI) and monitored virus titer in the supernatant by plaque assay. Two days after inoculation about 10^6^ infectious units/ml were produced (Fig. **2A**). Strikingly, RNAseq analyses indicated that viral transcripts account for about 88% of the total cellular transcriptome (Fig. **2B**, left panel). At the peak of SARS-CoV-2 infections of hiPSC-CM lines in two other studies, viral transcripts accounted for about 56% and 35% of the total cellular transcriptome (Fig. **2B**, center and right panel).

**Fig. 2:**
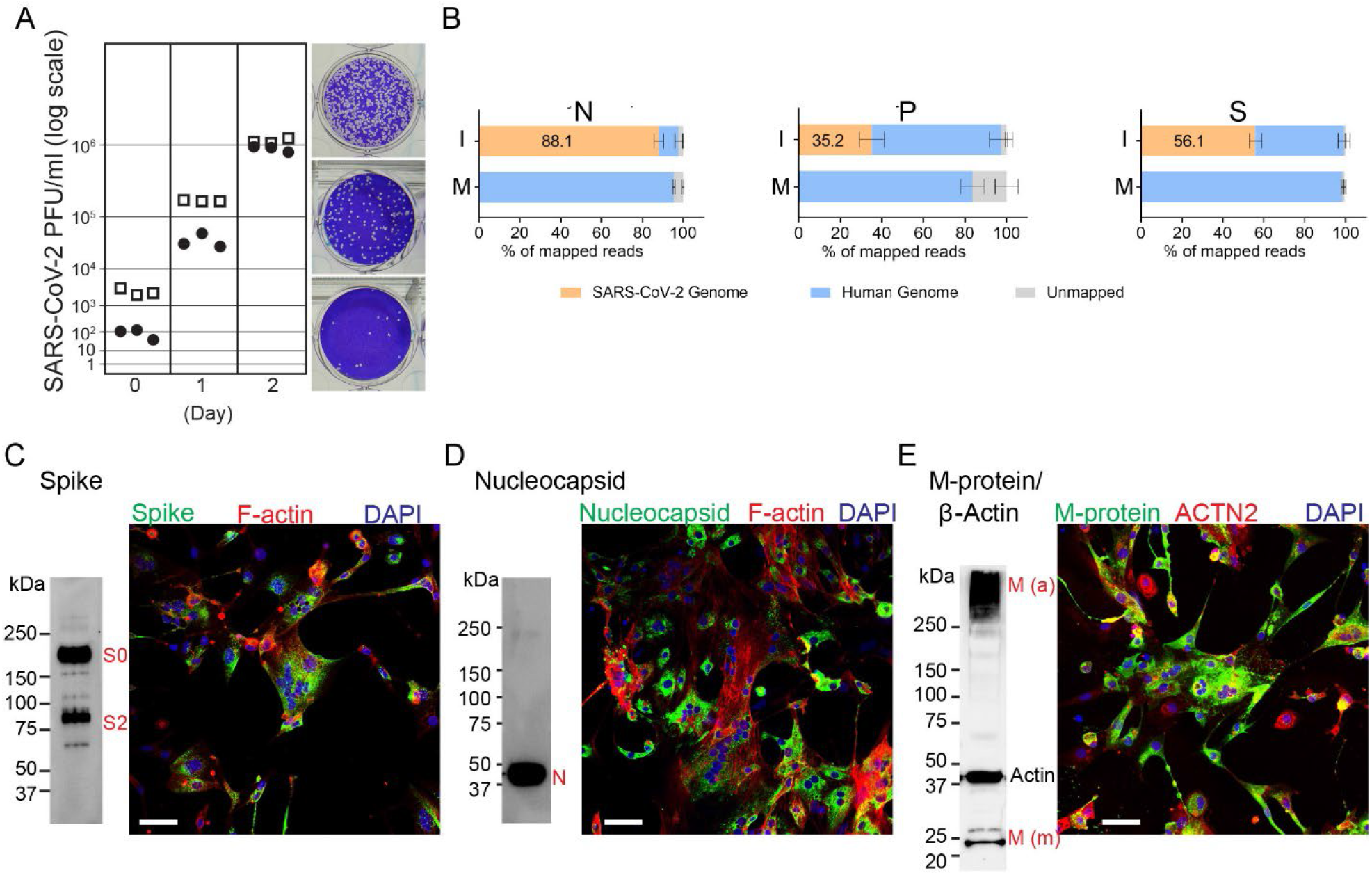
Efficient SARS-CoV-2 infection of hiPSC-CMs. (**A**) SARS-CoV-2 titers in two hiPSC-CM cell lines: open squares, hiPSC-CM#1; filled dots, hiPSC-CM#2, each data point represents one biological replicate. (**B**) Quantification of viral transcripts in infected hiPSC-CMs from this study (N) and two published studies: P=Perez-Bermejo et al., 2021 [21]; S=Sharma et al., 2020 [19]. I=infected cardiomyocytes; M=mock infected cardiomyocytes. (**C-E**) Companion immunoblots (left) and low-power IF confocal microscopy (right) of (**C**) SARS-CoV-2 spike glycoprotein (S0, S2), (**D**) nucleocapsid, (N) and (**E**) membrane (M) protein, monomer (m) and insoluble aggregate in hiPSC-CMs, 48 hours post-infection. Scale bar, 50 µm.

We then sought to document expression, processing and localization of the viral proteins. Immunoblot analyses of viral spike (S), nucleocapsid (N), and membrane (M) proteins confirmed high expression levels and accurate processing (Fig. **2C-E**, left panels). Immunofluorescence (IF) microscopy confirmed localization of all three proteins to the expected subcellular compartments (Fig. **2C-E**, right panels). Taken together, these analyses confirmed highly productive infection of hiPSC-CMs by SARS-CoV-2.

### Abundant progeny virions in exocytic vesicles

We then assessed by transmission electron microscopy (TEM) whether SARS-CoV-2 infection of hiPSC-CM recapitulates features characteristic of other coronavirus infections. TEM analyses revealed canonical double-membrane vesicles, endoplasmic reticulum-Golgi intermediate complex and smooth-walled exocytic vesicles containing numerous 65-90 nm particles (Fig. **3A**, yellow box). These are progeny virions with typical helical ribonucleocapsids surrounded by a membrane (Fig. **3A**, inset). Other characteristic features of coronavirus infections detected in hiPSC-CM include clustered membranes (Fig. **3B**, yellow arrows), vesicle packets filled with virus particles (Fig. **3C**, blue arrows) and exocytic vesicles filled with virus particles (Fig. **3D**, white arrows). Thus, TEM analyses of infected hiPSC-CM detected alterations of the cellular secretory pathway characteristic of coronavirus infections.

**Fig. 3:**
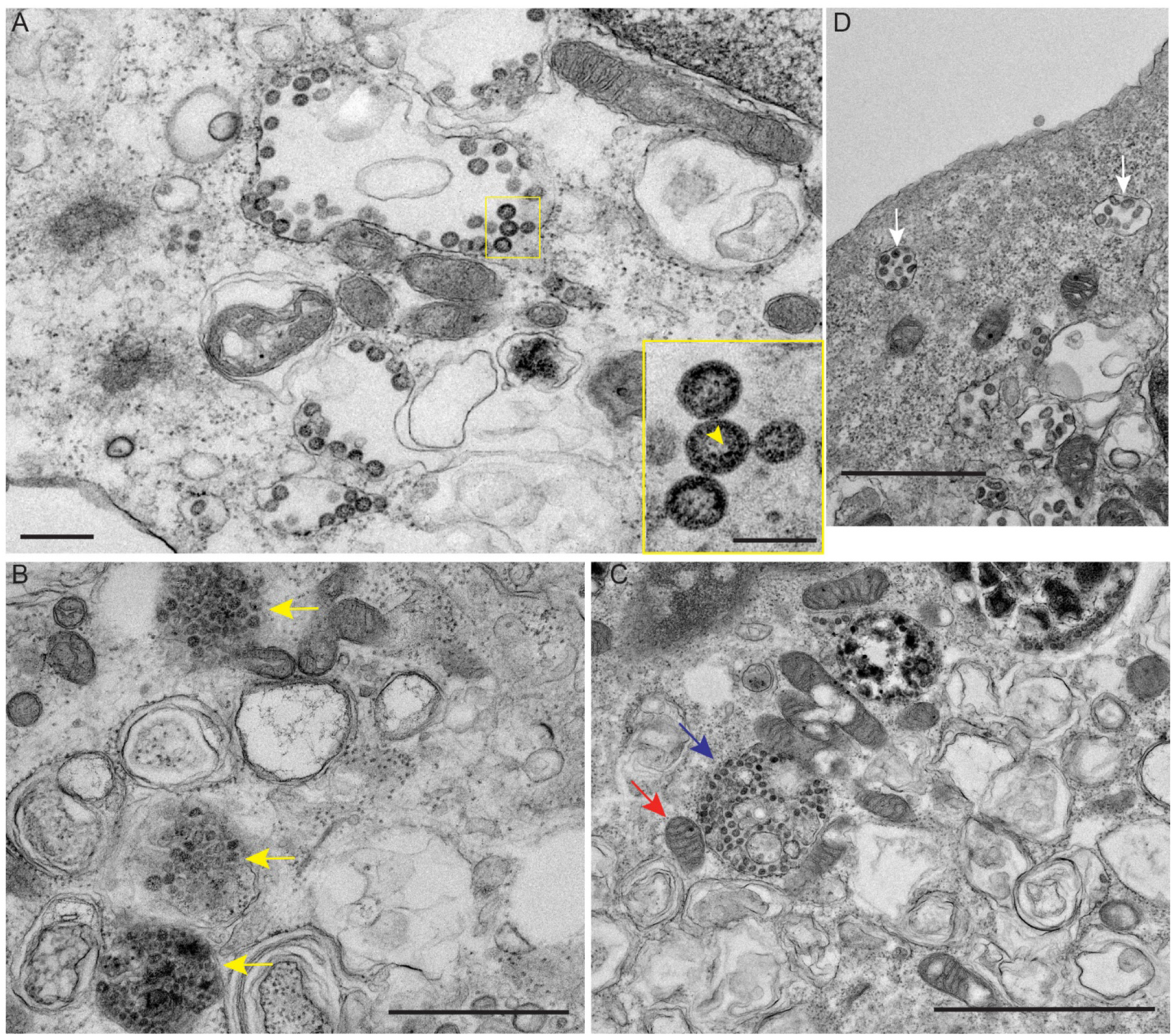
Maturation of SARS-CoV-2 in hiPSC-CMs. (**A**) TEM of SARS-CoV-2 infected hiPSC-CMs, 48 hours post-inoculation revealing numerous SARS-CoV-2 particles concentrated within vesicles (yellow box). Scale bar, 400 nm. Inset panel is high magnification of the viral particles, demonstrating electron-dense ribonucleocapsid structures (yellow arrow). Scale bar, 100 nm. (**B**) SARS-CoV-2 clustered membranes (yellow arrows). Scale bar, 1 µm. (**C**) SARS-CoV-2 vesicle packet (blue arrow); mitochondria (red arrow). Scale bar, 2 µm. (**D**) SARS-CoV-2 exocytic vesicles (white arrows). Scale bar, 1 µm.

### Virus-like particles with knob-like spikes on the cardiomyocyte surface

We assessed whether typical SARS-CoV-2 particles are present on the surface of hiPSC-CM by scanning electron microscopy (SEM), which revealed numerous particles on the plasma membrane. Fig. **4A** shows an hiPSC-CM heavily carpeted with SARS-CoV-2 particles (rightmost cell) contacting two less heavily carpeted hiPSC-CMs at upper and lower left with boundaries clearly demarcated, creating a patchwork mosaic. The inset magnifies the boundary highlighted by the yellow box. Viral particles cover the entire surface of the hiPSC-CM, including pseudo and filopodia and show typical knob-like spikes (Fig. **4B**). Thus, SEM analyses detected abundant *bona fide* virus-like particles on the hiPSC-CM surface, and individual cells produce different amount of virus particles.

**Fig. 4:**
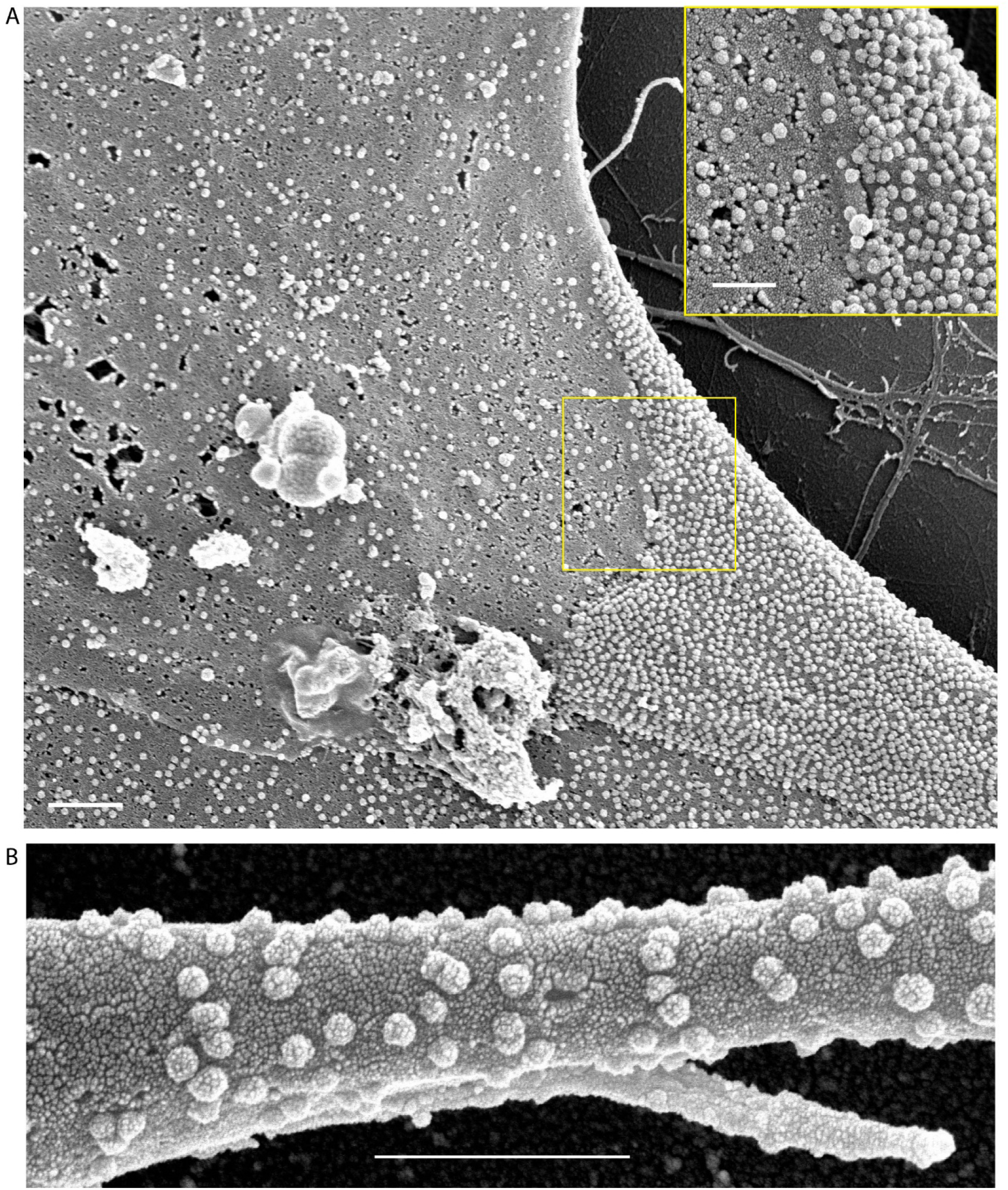
Surface of hiPSC-CMs infected with SARS-CoV-2. (**A**) Scale bar, 1 µm. Inset shows high magnification of the surface region within the yellow box. Scale bar, 500 nm. (**B**) High magnification SEM of hiPSC-CM filopodia dotted with SARS-CoV-2 viral particles. Scale bar, 1 µm.

### Cytopathic effects and fusion of infected cardiomyocytes

We also monitored the cytopathic effects of SARS-CoV-2 infection of hiPSC-CMs by IF confocal microscopy. In Fig. **5A** the nuclei of infected cells were stained with DAPI (blue), and the viral M protein and cytoskeletal alpha-actinin with specific antibodies (green and red, respectively). Fig. **5B** shows the same analyses on control uninfected hiPSC-CMs. In Fig. **5A** giant cells with central clusters of nuclei, named syncytia, were documented. These M-protein positive hiPSC-CMs demonstrated sarcomeric disassembly/fragmentation shown by disintegration of α-actinin Z-discs into randomly distributed puncta (Fig. **5C**). Neither syncytia formation nor cytoskeletal disassembly were observed in mock-infected cells (Fig. **5D**).

**Fig. 5:**
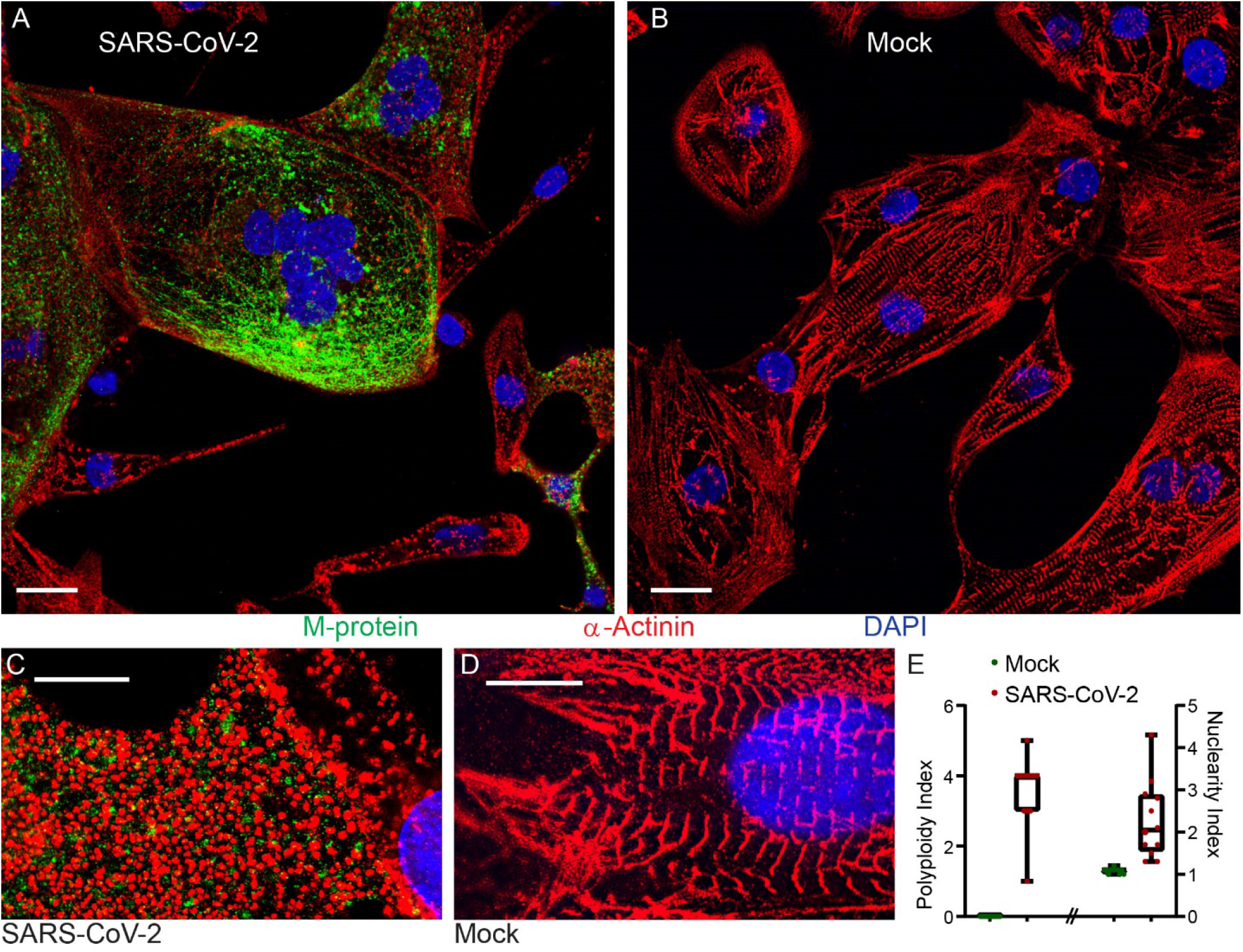
Cytopathic effects of SARS-CoV-2 in hiPSC-CMs. (**A-B**) IF confocal microscopy of SARS-CoV-2 infected (48 hours post-infection) or mock-infected hiPSC-CMs, respectively. Scale bar, 20 µm. (**C-D**) IF super resolution confocal microscopy of infected and mock-infected hiPSC-CMs, respectively. Scale bars, 10 µm. (**E**) Quantification of cell fusion in SARS-CoV-2 infected and mock infected hiPSC-CMs 48 hours post-inoculation. Polyploidy index is the average number of syncytia per field (n= 3 biological replicates). Nuclearity index is the average number of nuclei per cell per field (n= 3 biological replicates, p = 0.0002, two-tailed t-test). Box and whisker plots show median, upper and lower quartile, and extremes. Each data point represents one image field containing about 30 cells. Twelve image fields were counted per condition.

To quantify SARS-CoV-2 mediated hiPSC-CM fusion, α-actinin and SARS-CoV-2 M protein co-labeled cells were imaged by IF confocal microscopy and syncytia were counted. While no syncytia were observed for mock infected cells, ∼4 were counted per field of SARS-CoV-2 infected cells (Fig. **5E**, polyploidy index). As an alternative method to quantify fusion, we counted the number of nuclei per cell, finding an average of about 2 in infected cells, double that counted in the mock control (Fig. **5E**, nuclearity index). Thus, some infected cardiomyocytes fuse and cytoskeletal disintegration may favor syncytia formation.

### A fluorescent viral spike protein fuses cardiomyocytes

To characterize the mechanism of cell fusion, we engineered a SARS-CoV-2 full-length recombinant spike protein fused to modified Emerald green fluorescent protein at its carboxyl-terminus (CoV-2 S-mEm) (Fig. **6A**, left panel). We validated this reagent in Vero cells that, like hiPSC-CMs, express ACE2 but not TMPRSS2. In these cells CoV-2 S-mEm was appropriately cleaved (Fig. **6A**, right panel). Super resolution confocal microscopy localized CoV-2 S-mEm to hair-like plasma membrane extensions (Fig. **6B**). Fluorescent activated cell sorting confirmed CoV-2 S-mEm cell surface expression (Fig. **6C**). Live cell imaging tracked spread of CoV-2 S-mEm signal from cell to cell through membrane fusion, generating syncytia (Fig. **6D** and **S2** Movie).

**Fig. 6:**
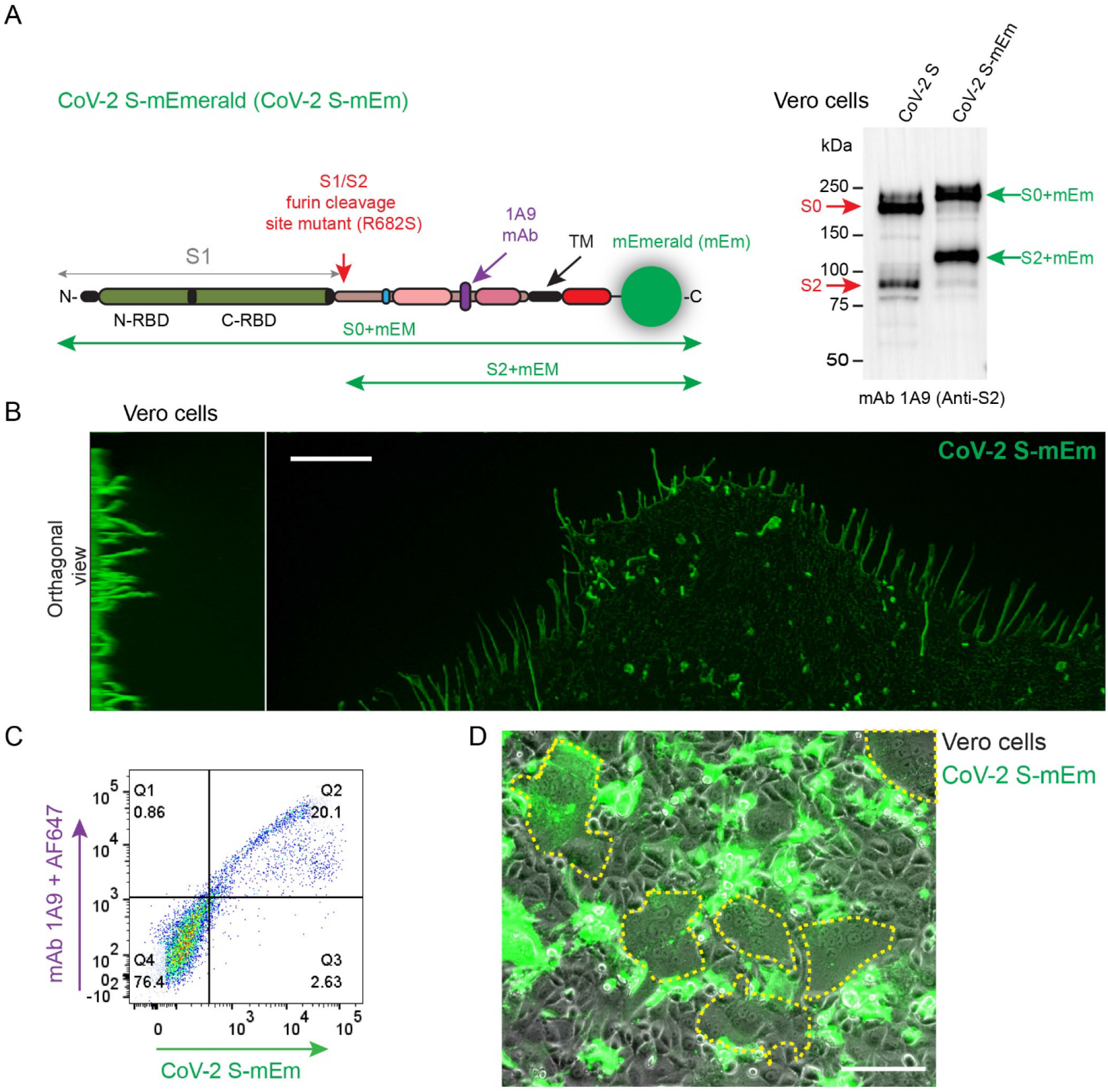
Expression and function of SARS CoV-2 spike protein tagged with mEmerald. (**A**) Left panel: schematic of SARS-CoV-2 S tagged with mEmerald (mEm) at the cytoplasmic tail. Cleavage at the S1/S2 furin site primes the spike protein for activation. S1, S1 subunit; S2, S2 subunit; N-/C-RBD, N-/C-terminal receptor binding domains; TM, trans-membrane segment. The fusion peptide is shown in blue and heptad repeat 1 and 2 in magenta and dark magenta, respectively. The location of the furin cleavage mutant, R682S is indicated. The monoclonal antibody 1A9, which was used to detect the spike proteins, binds to an exposed loop (purple) located close to heptad repeat 2. Right panel: immunoblot of the CoV-2 S and CoV-2 S-mEm proteins detecting their S0 and S2 subunits. (**B**) Super resolution confocal microscopy of CoV-2 S-mEM localization to Vero cell filopodia. Scale bar, 5 µm. (**C**) Cellular localization of the tagged spike protein in non-permeabilized HeLa cells transfected with the expression plasmid for S-mEm. This protein was detected either by fluorescence emission (horizontal axis) or by using spike-specific-mAb 1A9 and AF647 conjugated secondary-antibody (vertical axis). (**D**) Syncytia in Vero cells transfected with CoV-2 S-mEm are indicated by a dotted yellow line. Scale bar, 50 µm.

We then assessed whether CoV-2 S-mEm fuses cardiomyocytes. Despite overall transfection efficiency <5%, CoV-2 S-mEM expressing hiPSC-CMs produced syncytia with nuclei frequently arranged in clusters or rosettes (Fig. **7A**, and **S3** Movie). Some syncytia were characterized by circular or oval enucleated cytoskeletal “corpses” shown by F-actin phalloidin staining (Fig. **7B**, yellow arrows). Super resolution confocal microscopy demonstrated fluorescent signal at the tips of dynamic pseudo- and filopodia contacting neighboring hiPSC-CMs (Fig. **7C**, circle). Since hiPSC-CMs do not express TMPRSS2, we conclude that in these cells another protease must activate the SARS-CoV-2 spike.

**Fig. 7:**
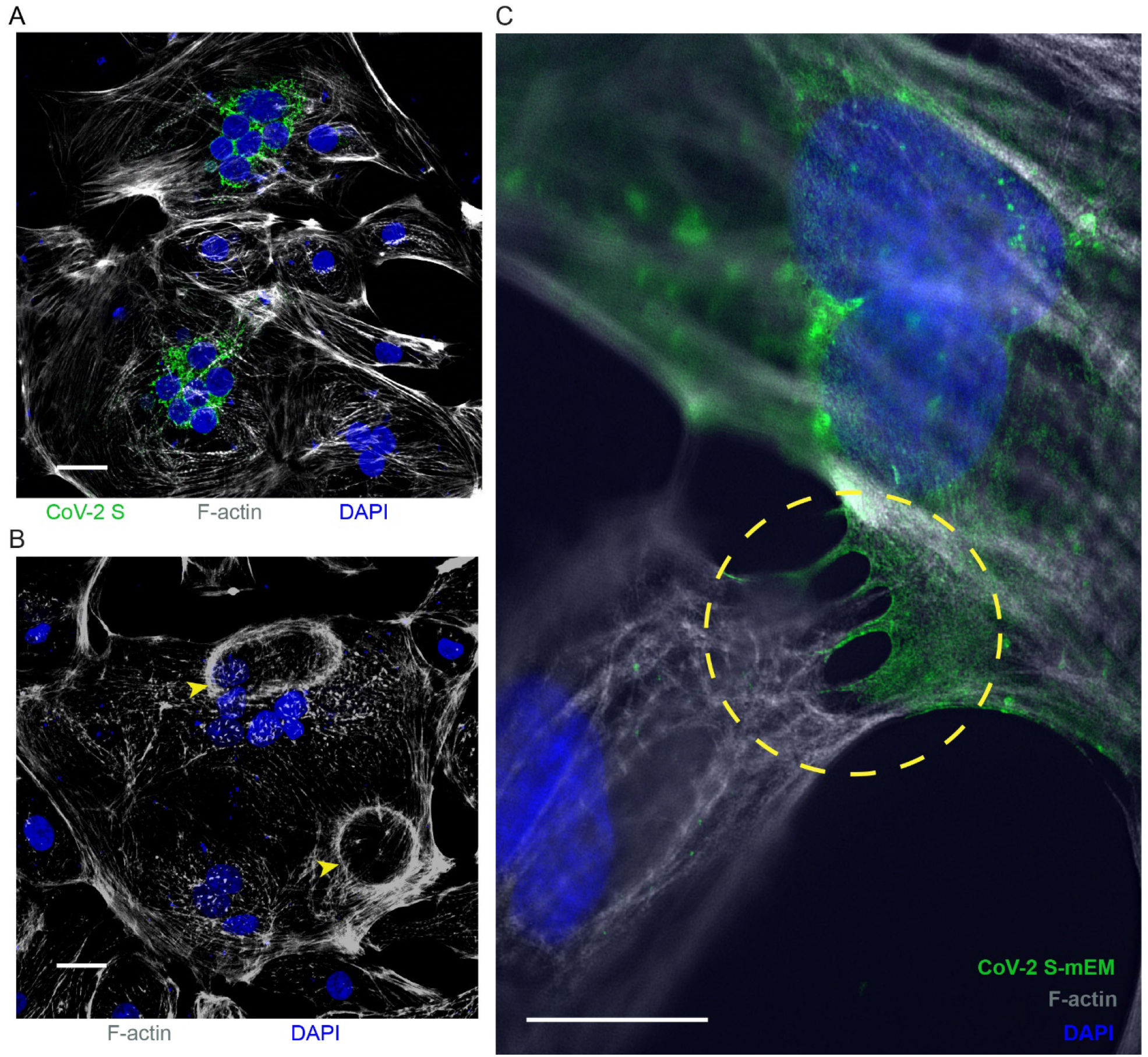
SARS-CoV-2 spike protein induces syncytia in hiPSC-CMs. (**A**) IF confocal microscopy of SARS-CoV-2 spike (CoV-2 S)-expressing hiPSC-CMs. Scale bar, 50 µm. Viral and cellular components visualized are indicated with their corresponding color below each panel. (**B**) IF confocal microscopy of CoV-2 S-expressing hiPSC-CM with enucleated actin cytoskeletal “corpses” (yellow arrows). (**C**) Super resolution confocal microscopy of CoV-2 S-mEM localization to hiPSC-CM filopodia directly contacting the sarcolemma of an adjacent hiPSC-CM (yellow circle). Scale bar, 2 µm.

### Furin activation of spike is required for cardiomyocyte fusion

Knowing that furin, a protease located in the *trans*-Golgi apparatus that contributes to SARS-CoV-2 spike activation, is expressed in hiPSC-CM (Fig. **1B**), we sought to block its function biochemically and genetically. For biochemical interference we used Decanoyl-RVKR-CMK (furin inhibitor, FI), a cell-permeable peptide-based molecule that irreversibly blocks its catalytic site. For genetic interference, we generated an expression vector differing from CoV-2 S through the single amino acid change R682S expected to inactivate the furin cleavage site [25] (Fig. **6A**, left panel).

We validated these approaches in Vero cells. The left panel of Fig. **8A** documents progressive inhibition of CoV-2 S protein processing (S0 cleavage into S1 and S2) by increasing concentrations of FI. The second and third panels show that fusion occurs in cells expressing CoV-2 S in the absence of FI, but not in its presence. The last panel shows that the R682S mutant of CoV-2 S is fusion-inactive.

**Fig. 8:**
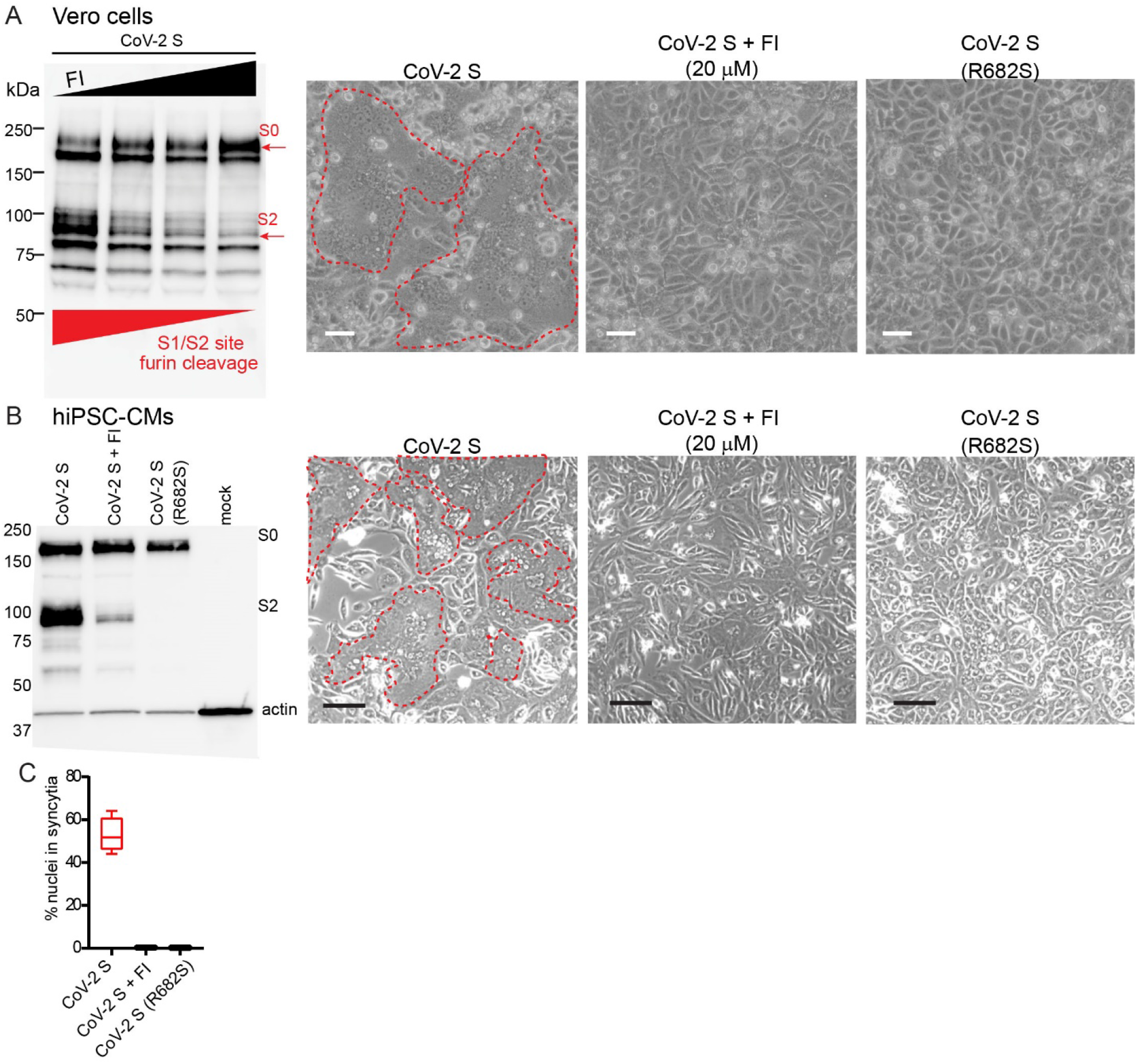
SARS-CoV-2 spike generated syncytia are blocked by a furin inhibitor or a furin-cleavage mutant. (**A**) (left panel) immunoblot analysis of CoV-2 S protein processing (S0 cleavage into S1 and S2) in Vero cells treated with increasing concentrations of FI (0 μM, 5 μM, 10 μM and 20 μM); cell lysates were separated by 4-15% SDS-PAGE under reducing conditions. (2^nd^ to 4^th^ panels) phase contrast images of Vero cells expressing CoV-2 S in the absence or presence of 20 µM FI, or of Vero cells expressing the R682S cleavage mutant, respectively, 72-hours after transfection. Syncytia are demarcated by a dashed broken red line. Scale bar, 100μm. (**B**) (left panel) gel analysis of CoV-2 S protein processing in the absence or presence of FI (20 µM), and of the processing of the CoV-2 S R682s furin cleavage mutant (R682S). Lysates of hiPSC-CM were separated by 4-15% SDS-PAGE under reducing conditions. (2^nd^ to 4^th^ panels) phase contrast images of hiPSC-CM expressing CoV-2 S in the absence or presence of 20 µM FI, or of Vero cells expressing the R682S cleavage mutant, respectively, 72-hours after transfection. Syncytia are demarcated by a dashed broken red line. Scale bar, 100μm. (**C**) Quantification of hiPSC-CM fusion. % nuclei in syncytia denotes the percent of total nuclei within syncytia 48 hr post transfection (n= 3 biological replicates, p <0.0001, ANOVA). Box and whisker plots for all quantification in this figure shows median, upper and lower quartile, and extremes.

Fig. **8B** shows that furin activation of spike is required also for cardiomyocyte fusion. The left panel documents strong inhibition of spike protein processing by high concentration of FI, and complete lack of processing of the R682S mutant. The other panels show that FI or the mutant inhibits fusion of cardiomyocytes. Fig. **8C** shows a quantitative analysis of hiPSC-CM fusion documenting approximately 99% inhibition by FI and by the mutation. Thus, expression of furin-activated SARS-CoV-2 spike protein in hiPSC-CM causes cell fusion that can be corrected pharmacologically.

### The MERS spike drives cardiomyocyte fusion with slow kinetics

Since SARS-CoV [26] and Middle East respiratory syndrome (MERS) [27] can cause cardiovascular complications, we asked whether their spike proteins can fuse hiPSC-CM. As negative control we used the spike protein of the common cold coronavirus 229E. Fig. **9A** shows correct processing of the MERS spike protein, and Fig. **9B-C** demonstrates that this protein induces syncytia formation. Comparative analyses indicated that the MERS spike protein drove syncytia production with slower kinetics than the SARS-CoV-2 spike, while the spike proteins of SARS-CoV and of the common cold coronavirus HCoV-229E were inactive. These levels of fusion activity correlate with the amounts of cardiovascular complications observed in infections with the respective viruses.

**Fig. 9:**
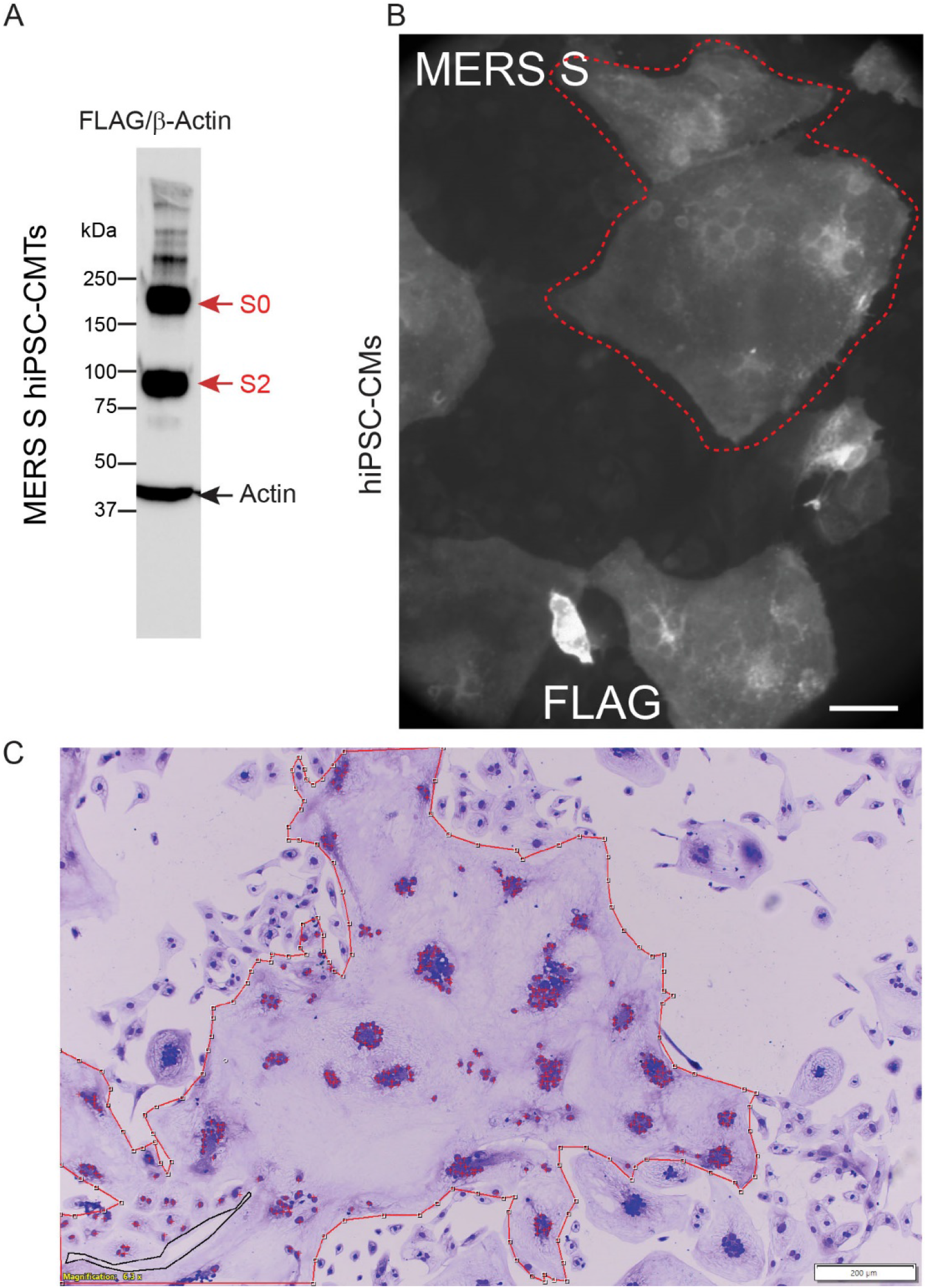
MERS spike-mediated syncytia. (**A**) Immunoblot of hiPSC-CM expressing recombinant MERS spike protein showing processing. The MERS spike S0 precursor and S2 cleaved subunit are detected through a FLAG epitope fused to the C-terminus. High molecular weight (>250 kDa) oligomers, including trimers, are detected. (**B**) Anti-FLAG IF microscopy of MERS spike protein-mediated hiPSC syncytia, largest example circled in red. Scale bar, 50 µm. (**C**) Bright field microscopy of crystal violet-stained hiPSC-CM expressing recombinant MERS spike protein at 5 days post-transfection. A composite syncytium is circled in red. Scale bar, 200 µm.

## Discussion

Viruses can cause cardiomyopathies, but the mechanisms of disease are difficult to characterize experimentally [28, 29]. The cardiac complications frequently observed in COVID-19 patients are usually tentatively attributed to aberrant host responses to acute respiratory infection. However, SARS-CoV-2 replication has occasionally been confirmed in autopsy cardiac tissue. While animal models to study SARS-CoV-2 infections of the heart are being developed, we have characterized virus spread in hiPSC-CM. Infection of these cells was highly efficient, with the virus taking over almost 90% of the cellular transcriptome. SARS-CoV-2 infection re-shaped subcellular morphologies [30, 31], secretory vesicles were filled with viral progeny, and virus particles with knob-like spikes carpeted the cardiomyocyte surface.

Human iPSC-CM are permissive to SARS-CoV-2 infection. Two previous studies documented that viral transcript account for up to 35% or 55% of the hiPSC-CM transcriptome, respectively [19, 21], comparing favorably but not exceeding 65% cellular transcriptome takeover reported after infection of Vero cells [32]. We do not know why cell takeover was more extensive in our hiPSC-CM infections, but we note that the expression levels of the ACE2 receptor transcript are high in our study, intermediate in that of Sharma et al., and low in that of Perez-Bermejo. Thus, levels of ACE2 expression correlate with levels of cellular transcriptome takeover.

We document here syncytia formation in infected hiPSC-CM. Most spike protein produced in infected cells engages the other viral components early in the secretory pathway, and progeny virus particles bud intracellularly. Nevertheless, some spike protein reaches the surface, where it can interact with the ACE2 receptor on the surface of other cells, starting cell-cell fusion. Indeed, cells infected by SARS-CoV-2 can fuse with neighboring cells to form large multinucleated syncytia, which have been documented in autopsy material from the respiratory tract of COVID-19 patients [33-35] and in *in vitro* models of airway epithelia infection [36-38] [39]. However, cell-cell fusion was not reported in previously published hiPSC-CM studies. In our studies, expression of ACE2 receptor was higher and infection more efficient. Thus, higher expression levels of both the spike protein and its receptor may account for more pronounced cell fusion in our study.

Proper proteolytic activation of the viral spike is required both for cell entry and cell-cell fusion [40]. In airway epithelial cells, fusion is triggered by the protease TMPRSS2, which processes the spike protein to set free its fusion peptide and elicit membrane fusion. SARS-CoV-2 has evolved a multibasic site at the S1-S2 boundary that allows for proteolytic processing of spike by furin in the *trans*-Golgi complex of the producer cell, rather than during entry into target cells. In 293T and in Vero cells furin is not absolutely required for cell-cell fusion, while the multibasic site and the concomitant presence of TMPRSS2 sustain this process [41]. Since TMPRSS2 is not expressed in cardiomyocytes, in these cells another protease may trigger fusion. This suggestion is consistent with the observation that coronaviruses can recruit a cohort of different protease to confer fusion competence to their spike [42, 43]. Thus, in the heart and other extrapulmonary organs SARS-CoV-2 pathogenesis may be independent of TMPRSS2. Understanding the mechanisms of SARS-CoV-2 spread in hiPSC-CM can inform antiviral therapies. We show here that the peptide-based Decanoyl-RVKR-CMK furin protease inhibitor abolishes cytopathology in cardiomyocytes. A derivative of this furin inhibitor with improved bioavailability and specificity could be advanced towards clinical trials as an inhibitor of COVID-19 induced cardiac disease.

## MATERIALS AND METHODS

### Spinner culture cardiac differentiation of human-iPSCs

Obtained under Mayo Clinic IRB protocol, patient and control human fibroblast-derived iPSCs were maintained in mTESR1 basal media with mTESR supplement on plates coated with Geltrex (in DMEM/F12 media). Undifferentiated hiPSCs were transitioned and expanded in suspension/spinner culture in DMEM/F-12 plus Glutamax, StemPro supplement, BSA and bFGF with Rock Inhibitor Y27632 combined with mTESR1 media, and then chemically differentiated by CHIR/IWP-4 into CMs in RPMI 1640 plus B27 minus insulin supplement as beating aggregates. Detailed spinner culture cardiac differentiation protocol is available from J.W.S. upon request. Differentiated hiPSC-CMs were maintained in Gibco^™^ Cardiomyocyte Maintenance Medium and attached to fibronectin-coated glass coverslips. Human H9 embryonic stem cells (WiCell) were chemically differentiated into CMs using an analogous protocol in monolayer culture. EdU (5-ethynyl-2’-deoxyuridine) labeling of growing hiPSC-CMs and detection were done as described by the manufacturer (Thermo-Fisher).

### SARS-CoV-2 infection of hiPSC-CM

SARS-CoV-2/UW-001/Human/2020/Wisconsin (UW-001) was isolated from a mild case in February 2020 and passaged in VeroE6 cells expressing TMPRSS2. The virus was used to infect hiPSC-CMs in monolayer at MOI of 1.0 to 0.001 for 30 minutes at 37ºC. Unbound virus was then washed-off and fresh media replaced. At the various time points, cells were fixed or extracted, and samples were collected, and the vessels decontaminated. An MOI of 0.01 for 24-48 hours proved optimal for observing early stages of SARS-CoV-2 infection in hiPSC-CMs. Beyond 72 hours, even at low starting MOI, cytopathic lysis overwhelmed hiPSC-CM cultures. Highly permissive SARS-CoV-2 infection was observed in 3 different, equivalently differentiated hiPSC-CMs from unrelated donors.

### Virus titration

SARS-CoV-2 infectious virus produced by hiPSC-CM was titered by plaque-forming assay done in confluent Vero E6/TMPRSS2 cells. 12-wells tissue culture plates were infected with supernatant (undiluted and 10-fold dilutions from 10 to 10^5^) for 30 minutes at 37°C. After initial exposure, the Vero/TMPRSS2 cells were washed three times to remove unbound virus and the media was replaced with 1.0% methylcellulose-media. After an incubation of three days at 37°C, the cells were fixed and stained with crystal violet solution and plaque number counted to determine plaque-forming units (PFU)/ml.

### RNA sequencing

The hiPSC CMs were infected with SARS-CoV-2/UW001/Human/2020/Wisconsin (UW-001) at MOI of 0.01. Cells were lysed in Trisol and were kept at -80°C. Total RNA of the lysate was extracted using Direct-zol RNA Miniprep kit (R2050). Library preparation and sequencing was performed at Mayo Clinic Genome Analysis Core (GAC).

Briefly, cDNA libraries were prepared using 100 ng of total RNA according to the manufacturer’s instructions for the Illumina TruSeq Stranded Total RNA Sample Prep Kit with Ribo-Zero Gold (Illumina, San Diego, CA). The concentration and size distribution of the completed libraries were determined using an Agilent Bioanalyzer DNA 1000 chip (Santa Clara, CA) and Qubit fluorometry (Invitrogen, Carlsbad, CA). Libraries were sequenced at three samples per lane to generate approximately 119 to 137 million fragment reads per sample following Illumina’s standard protocol using the Illumina cBot and HiSeq 3000/4000 PE Cluster Kit. The flow cells were sequenced as 100 × 2 paired end reads on an Illumina HiSeq 4000 using HiSeq 3000/4000 sequencing kit and HD 3.4.0.38 collection software. Base-calling was performed using Illumina’s RTA version 2.7.7.

### Bioinformatics and data analysis

The quality of the raw RNA-seq data was assessed by fastqp v0.20.1 [44], and quality reads were filtered and aligned against human genome (hg19) using STAR alignment (v2.7.8a) [45] in galaxy platform (https://usegalaxy.org). The aligned reads were counted using htseq-count v0.9.1 [46] and 0.5 read counts per million (CPM) in at least two samples was used as an expression threshold. Trimmed mean of M values normalized (TMM) [47] and log2 transformed data was used for plotting heatmaps and differential analysis in limma [48]. For the detection of viral transcripts, quality filtered reads were aligned against SARS-CoV-2 genome (MT039887.1) using BWA-MEM v0.7.17.1 (https://arxiv.org/abs/1303.3997). Alignment summary statistics was computed using samtools idxstats v2.0.3 [49]. The same workflow was used to re-analyze the data of ([19], GSE150392) and ([21], GSE156754) RNA-seq data except for SARS-CoV-2 genome (MN985325.1). The raw RNA-seq data from this study are available at Gene Expression Omnibus with accession number xxxx [to be added].

### Immunocytochemistry

Coverslips were fixed with neutral buffered formalin for 15 min at room temperature, washed with PBS/0.05% Tween-20 and blocked in (PBS/5% normal goat serum or 3% BSA/0.3% Triton X-100) at room temperature for 1 hour. Coverslips were incubated in primary antibodies diluted in (PBS/1%BSA/0.3% Triton X-100) overnight at 4°C, washed extensively and incubated with diluted secondary antibodies (1:400) at room temperature for 1 hour, then DAPI stained for 10 min at room temperature. Coverslips were mounted on slides with Prolong Gold Antifade Mountant (ThermoFisher) and stored at 4°C. Coverslips were imaged using a Zeiss LSM780 or Elyra PS.1 Super Resolution confocal microscope. Antibodies and reagents for immunocytochemistry included: ACTC1 (Actin α-sarcomeric mouse mAb clone 5C5 (Sigma), Phalloidin Alexa Fluor-568 conjugated (Invitrogen), SARS-CoV-2 Spike mAb clone 1A9 (GeneTex), SARS-CoV-2 M rabbit polyclonal Ab (Argio Biolaboratories), SARS-CoV-2 Nucleocapsid clone 1C7 (Bioss Antibodies), ACE2 goat polyclonal Ab (R&D Systems) and ATP2A2/SERCA2 rabbit polyclonal Ab (Cell Signaling).

### Transmission electron microscopy

Cells were washed with PBS and placed in Trump’s universal EM fixative[50] (4% formaldehyde, 1% glutaraldehyde in 0.1 M phosphate buffer, pH 7.2) for 1 hr or longer at 4º C. After 2 rinses in 0.1 M sodium phosphate buffer (pH 7.2), samples were placed in 1% osmium tetroxide in the same buffer for 1 hr at room temperature. Samples were rinsed 2 times in distilled water and dehydrated in an ethanolic series culminating in two changes of 100% acetone. Tissues were then placed in a mixture of Epon/Araldite epoxy resin and acetone (1:1) for 30 min, followed by 2 hrs in 100% resin with 2 changes. Finally, samples were placed in fresh 100% resin polymerized at 65º C for 12 hrs or longer. Ultrathin (70-90 nm) sections were cut with a diamond knife and stained with lead citrate. Images were captured with a Gatan digital camera on a JEOL 1400 plus transmission electron microscope operated at 80KeV.

### Scanning electron microscopy

Fixed in Trump’s (1% glutaraldehyde and 4% formaldehyde in 0.1 M phosphate buffer, pH 7.2) [50], tissue was then rinsed for 30 min in 2 changes of 0.1 M phosphate buffer, pH 7.2. Following dehydration in progressive concentrations of ethanol to 100% the samples were critical-point dried. Specimens were then mounted on aluminum stubs and sputter coated with gold/palladium. Images were captured on a Hitachi S4700 scanning electron microscope operating at 3kV.

### HeLa and Vero cells

HeLa cells were cultured in Dulbecco’s modified Eagle’s medium (DMEM) supplemented with 10% FBS. Vero-hSLAM (Vero cells stably expressing human signaling lymphocyte activation molecules, kindly provided by Y. Yanagi; these cells are described simply as Vero cells in this manuscript) [51] were maintained in DMEM supplemented with 10% FBS and 0.5 mg of G418/ml. All cell lines were incubated at 37°C with 5% CO_2_.

### Plasmids and mutagenesis

The codon-optimized SARS-CoV2 S-protein gene (YP_009724390) was synthesized by Genewiz in a pUC57-Amp plasmid (kindly provided by M. Barry). The S-protein coding sequence was cloned into a pCG mammalian expression plasmid [52] using unique restriction sites *Bam*HI and *Spe*I. The SARS CoV S-protein (VG40150-G-N) and the MERS S-protein (C-terminal FLAG tag, VG40069-CF) purchased from Sino Biological, were cloned into the pCG vector for comparative studies. The SARS-CoV-2 S-mEmerald construct was made by cloning the mEmerald sequence (Addgene, Plasmid #53976) to the C-terminal end of the SARS CoV-2 S-protein in the pCG expression vector. A flexible 6 amino acid-linker (TSGTGG) was used to separate the two proteins. All expression constructs were verified by sequencing the entire coding region. The R682S furin cleavage mutation was introduced into the SARS-CoV-2 S expression plasmid by QuikChange site-directed mutagenesis (Agilent Technologies, Santa Clara, CA) according to the manufacturer’s instructions. The clones were verified by sequencing the S-protein gene in the vicinity of the mutation. Two independent clones were tested.

### Immunoblots

Vero cells were transfected with spike protein expression constructs using the GeneJuice transfection reagent (Novagen). The indicated S-protein expression constructs (1 µg) were transfected into 2.5×10^5^ Vero cells in 12-well plates. Thirty-six hours post-transfection, extracts were prepared using cell lysis buffer (Cell Signaling Technology, #9803) supplemented with cOmplete protease inhibitor cocktail (Roche, Basel, Switzerland) and the proteins separated by sodium dodecyl sulfate-polyacrylamide gel electrophoresis (SDS-PAGE) (4 to 15% gradient) under reducing conditions. For hiPSC-CMs transfected with CoV-2 S (2 µg/well in 6-well plates), extracts were prepared in cell lysis buffer as above (but also including PMSF), and separated by SDS-PAGE under reducing (β-Mercaptoethanol) or non-reducing conditions. The S-proteins were visualized on an immunoblot using the anti-S specific monoclonal antibody 1A9 (GeneTex, GTX632604; 1:2000 dilution) which binds the S2 subunit of SARS CoV and SARS-CoV-2 S-proteins. An anti-mouse horseradish peroxidase (HRP)-conjugated secondary antibody was used to reveal the bands. MERS S-protein was detected using a monoclonal anti-FLAG M2-HRP conjugated antibody (SIGMA, A8592 @ 1:2000) which bound to a C-terminal FLAG-tag. The expression of the mEmerald tag was verified using a polyclonal anti-GFP antibody (Abcam, ab290 @ 1:5000). For hiPSC-CMs infected with SARS-CoV-2 (MOI 0.01, 48 hours), extracts were prepared in cell lysis buffer as above (but also including PMSF), separated by SDS-PAGE and blotted with S, M and N antibodies as described under Immunohistochemistry above.

### Cell-cell fusion assays

For spike glycoprotein-mediated cell-to-cell fusion, 1.5×10^5^ Vero cells in 24-well plates were transfected with 0.5 µg of the indicated S-protein expression vector using the GeneJuice transfection reagent (Novagen) and syncytia formation monitored for 24-48 hours post-transfection. Images were collected by Nikon Eclipse TE300 using NIS-Elements F 3.0 software (Nikon Instruments, Melville, NY, USA). For recombinant spike glycoprotein-mediated fusion in hiPSC-CMs, subconfluent day-20 differentiated cells plated on fibronectin-coated glass coverslips in 6-well plates were transfected with 1-2 µg plasmid using Lipofectamine 3000. For CoV-2 S-mEm in hiPSC-CM experiments syncytia formation became obvious within 6 hours of transfection.

### Furin inhibitor treatment

Furin Inhibitor I (Decanoyl-RVKR-CMK, Calbiochem, #344930) dissolved in DMSO was added to Vero or hiPSC-CM cell culture medium 2-hours post transfection. Cell-cell fusion was followed for 72-hours (for Vero cells) and 5 days for hiPSC-CMS with media and inhibitor refreshed on day-3. Whole cell extracts were separated on SDS-PAGE and immunoblotted for SARS-CoV-2 S as described above or cells fixed and stained by crystal violet.

### Quantification of hiPSC-CM fusion

Human iPSC-CM were plated on 35 mm round glass bottom dishes and transfected with SARS-CoV-2 spike protein, with or without furin inhibitor treatment, or transfected with SARS-CoV-2 R682S mutant spike protein, as previously described. Bright-field microscopy images were taken at 10x magnification from randomly chosen areas of each culture dish. Five images from three independent replicates were counted for each condition. Images were manually counted for number of nuclei, number of syncytia, and number of nuclei per cell using Olympus Dimension cellSens software. Percent nuclei within syncytium denotes the percent of total nuclei counted within syncytium at 48 hours post-transfection. Syncytia are defined by a cell containing three or more nuclei. ANOVA statistical analysis was carried out using GraphPad Prism software.

### Fluorescence-activated cell sorting

To determine S-protein cell surface expression levels, HeLa cells (8 × 10^5^ in a 6-well plate) were transfected with the indicated S-protein expression plasmids (2 µg using GeneJuice transfection reagent). Thirty-six hours post-transfection, cells were washed in PBS and detached by incubating with Versene (Life Technologies) at 37°C for 10 min. The resuspended cells were washed twice with cold fluorescence-activated cell sorter (FACS) wash buffer (phosphate buffered saline, 2% FBS, 0.1% sodium azide) and then incubated with the anti-S-protein mAb 1A9 (GeneTex; 1:50 dilution) for 1 hour on ice. Cells were washed three times with cold FACS wash buffer and incubated with an AF647-conjugated secondary antibody (Thermo Fisher Scientific, a21235 @ 1:200) for 1 hour on ice. After three washes with FACS wash buffer, cells were fixed in 4% paraformaldehyde and analyzed with a FACSCalibur (BD Biosciences, San Jose, CA) cytometer and FlowJo software (Tree Star Inc., Ashland, OR)

### Time lapse confocal microscopy

Vero cells were sparsely plated on a glass-bottom 35-mm dish and transfected with 1 µg of the SARS-CoV-2 S-mEmerald expression construct using GeneJuice transfection reagent. Time lapse confocal microscopy with images taken every 30-40 minutes for 12-hours, was performed 24-hours post-transfection on a Zeiss LSM780 equipped with a heated CO_2_ chamber. For time-lapse confocal fluorescence video microscopy of CoV-2 S-mEm spike glycoprotein transfected hiPSC-CMs, images were captured every 30 minutes over a 12-hour time period starting 24 hours after transfection on a Zeiss LSM780 equipped with a heated CO_2_ chamber.

## Acknowledgments

We thank Jeff Salisbury and the Mayo Clinic Microscopy and Cell Analysis Core facility for experimental and technical support; Tim L. Emmerzaal and Tamás Kozicz for experimental support; Mike Barry for the SARS-CoV-2 spike coding sequence; Peter Rottier, Mathieu Mateo, Christian Pfaller, Michael Muehlebach and Christoph Springfeld for insightful comments on the manuscript; and the Mayo Clinic Graduate School of Biomedical Sciences for graduate student support (D.R.P. and D.J.C.). We also than Andrew Badley and the Mayo Clinic COVID-19 Taskforce for support. Wanek Family Program for HLHS-Stem Cell Pipeline: Timothy J. Nelson (Director), Boyd Rasmussen and Frank J. Secreto.

## Supporting information

**S1 Table**. Affymetrix microarray analyses of ACE2 and TMPRSS2 expression in H9 human embryonic stem cells.

**S2 Movie**. Intercellular spread of CoV-2 S-mEm spike protein and development of Vero cell syncytia.

**S3 Movie**. Intercellular spread of CoV-2 S-mEm spike protein and development of hiPSC-CM syncytia.

**S1 Table.** ACE2 and TMPRSS2 expression in H9 human embryonic stem cells.

| Probe set | Day 0 | Day 8 | Day 20 | Day 50 |
| --- | --- | --- | --- | --- |
| *ACE2*<br>219962_at | 11.83 P** | 4.53 P | 462.73 P | 82.48 P |
| *ACE2*<br>222257_s_at | 6.61 A\$ | 16.74 P | 672.49 P | 119.02 P |
| *TMPRSS2*<br>1570433_at | 9.75 A | 15.73 A | 11.82 A | 17.11 A |
| *TMPRSS2*<br>205102_at | 79.07 A | 47.56 A | 104.44 A | 80.07 A |
| *CTSB*<br>213275_x_at | 355.25 P | 1330.85 P | 1841.63 P | 1599.04 P |

\* Affymetrix microarray numerical values across an individual probe set
+ P (present): transcript is significantly ( $P < 0.05$ ) expressed compared with perfectly matched and mismatched (background) probe sets
\$A (absent): transcript is not significantly ( $P > 0.05$ ) expressed

## Notes

### Competing Interest Statement

The authors have declared no competing interest.

